# MLL4 regulates palatal salivary gland development

**DOI:** 10.64898/2026.11.11.766049

**Authors:** Bruno Pasqua, Hunmin Jung, Yungki Park, Rose-Anne Romano, Soo-Kyung Lee, Jae W. Lee, Jung-Mi Lee, Hyuk-Jae Edward Kwon

## Abstract

Palatal glands are minor mucous salivary glands that contribute to oral lubrication and mucosal defense through the secretion of mucus and protective factors such as secretory immunoglobulin A. They are also the most common site of benign and malignant minor salivary gland tumors. Despite their physiological and clinical importance, the molecular mechanisms underlying palatal gland development remain poorly understood. Given the essential role of the epigenetic regulator MLL4 (KMT2D) in craniofacial development, we investigated its role in palatal gland development using a conditional knockout (cKO) mouse model with tissue-specific deletion of *Mll4* in the palatal mesenchyme (*Mll4^fl/fl^;Osr2-Cre*). *Mll4*-cKO mice exhibited palatal gland hypoplasia from birth through adulthood, accompanied by ectopic cartilage formation in the soft palate. RNA-seq analysis revealed downregulation of genes and biological processes associated with salivary gland development, branching morphogenesis, and FGF and WNT signaling, together with upregulation of biological processes related to chondrocyte proliferation, consistent with the glandular hypoplasia and ectopic cartilage phenotypes. *In situ* hybridization and immunofluorescence analyses confirmed reduced mesenchymal *Fgf10* expression, decreased epithelial SOX9 and *Edar* expression, and localized accumulation of SOX9-positive cells within the ectopic cartilage. Ki67 immunostaining demonstrated reduced epithelial cell proliferation without altered apoptosis in *Mll4*-cKO palatal glands. Collectively, these findings identify MLL4 as an essential epigenetic regulator of palatal gland morphogenesis that coordinates mesenchymal–epithelial signaling while maintaining normal developmental programs within the soft palate.

## Introduction

Palatal glands are minor salivary glands located within the submucosa of the hard and soft palate beneath stratified squamous epithelium (Dayan et al., 2000). They consist predominantly of mucous acini that produce viscous, mucin-rich saliva (Hakami & Hand, 2018; Nakamura et al., 2001). Unlike the major salivary glands, palatal glands possess relatively short excretory ducts that open directly into the oral cavity (Okamoto et al., 2008). Although minor salivary glands produce substantially less saliva than the major salivary glands, they contribute significantly to oral homeostasis by providing a large proportion of the mucus and secretory immunoglobulin A that lubricate and protect the oral mucosa (Crawford et al., 1975; Dawes & Wood, 1973). Palatal glands are also the most common site of minor salivary gland neoplasms (Aluffi Valletti et al., 2024; Weisberger et al., 1979). Despite this physiological and clinical importance, the development of the palatal glands has received comparatively little attention.

During development, palatal glands arise as epithelial invaginations extending from the palatal epithelium into the underlying mesenchyme, followed by branching morphogenesis, lumenization, and acinar differentiation (Nielsen & Westergaard, 1971; Shinzato et al., 2004). Rodent studies have primarily described the histological maturation, ultrastructural organization, and secretory composition of palatal glands (Hakami & Hand, 2018; Hakami et al., 2014; Nakamura et al., 2001; Okamoto et al., 2008), whereas the molecular mechanisms regulating their morphogenesis remain poorly understood. In contrast, the development of the major salivary glands has been studied extensively and is driven by reciprocal epithelial–mesenchymal interactions, which coordinate epithelial bud initiation, progenitor cell expansion, branching morphogenesis, and terminal differentiation (Patel & Hoffman, 2014). Mesenchyme-derived FGF10 signals through epithelial FGFR2b to promote epithelial progenitor proliferation and branching morphogenesis (Chibly et al., 2022; Lombaert et al., 2013; Steinberg et al., 2005). Downstream of FGF10, SOX9 establishes and maintains the distal epithelial progenitor population required for branching morphogenesis (Chatzeli et al., 2017), while additional signaling pathways, including EDAR and WNT, further contribute to glandular morphogenesis (Häärä et al., 2011; Patel et al., 2011). Because palatal glands similarly develop through epithelial invagination into neural crest-derived palatal mesenchyme, mesenchymal signaling is expected to play a fundamental role in their morphogenesis. Whether similar epithelial–mesenchymal signaling mechanisms regulate palatal gland development remains unknown.

The histone methyltransferase MLL4 (mixed-lineage leukemia 4; also known as KMT2D) is an epigenetic regulator that catalyzes monomethylation and dimethylation of histone H3 lysine 4 (H3K4me1/2), which are modifications associated with enhancer activation and cell type-specific gene expression during development (Froimchuk et al., 2017; Xie et al., 2023). Pathogenic variants in *KMT2D* cause Kabuki syndrome, a congenital disorder characterized by distinctive craniofacial features, skeletal abnormalities, and developmental delay (Boniel et al., 2021). Although craniofacial abnormalities are well recognized in Kabuki syndrome, the developmental role of MLL4 in the palatal glands has not been investigated. Consistent with its broader role in craniofacial development, MLL4 is required for normal palate development, and our previous work using the same palatal mesenchymal *Mll4* deletion model demonstrated an essential role for MLL4 in postnatal palate growth and midpalatal suture development (Lee et al., 2025; Shpargel et al., 2020). Based on these findings, we hypothesized that mesenchymal MLL4 also regulates palatal gland development, potentially through epithelial–mesenchymal interactions.

In this study, we investigated the role of mesenchymal MLL4 in palatal gland development using *Mll4^fl/fl^;Osr2-Cre* mice. Palatal mesenchyme-specific deletion of *Mll4* resulted in palatal gland hypoplasia and ectopic cartilage formation in the soft palate. RNA-seq analysis revealed dysregulation of biological processes associated with FGF signaling, cartilage development, and chondrocyte proliferation, consistent with the observed phenotypes. *In situ* hybridization and immunofluorescence confirmed reduced mesenchymal *Fgf10* expression, together with decreased epithelial SOX9 and *Edar* expression and reduced epithelial proliferation. These findings reveal an essential role for mesenchymal MLL4 both in coordinating epithelial–mesenchymal signaling during palatal gland morphogenesis and in preventing ectopic cartilage formation in the soft palate, and support a non-cell-autonomous mechanism in which mesenchymal MLL4 is required for normal palatal gland development.

## Materials and methods

### Mouse strains

*Osr2-Cre* (Chen et al., 2009) and *Mll4^fl/fl^* (Lee et al., 2013) mouse lines were used to generate a palatal mesenchyme-specific *Mll4-*cKO model (Lee et al., 2025). Experimental *Mll4*-cKO mice and littermate controls were generated by crossing *Mll4^fl/+^;Osr2-Cre* males with *Mll4^fl/fl^* females, and the colony was maintained on a C57BL/6J background. The *ROSA26^mT/mG^* reporter line (denoted *mTmG*) was crossed with the *Osr2-Cre* line to label *Osr2-Cre*-active cells. All animal procedures were approved by the Institutional Animal Care and Use Committee at the University at Buffalo.

### Histology and immunofluorescence staining

Mice were euthanized at embryonic days 17.5 and 18.5 (E17.5 and E18.5), at birth (P0.5), and at postnatal stages (1, 3, 6, and 10 weeks). Head tissue specimens were fixed in 4% paraformaldehyde, dehydrated through a graded ethanol series, embedded in paraffin, and sectioned at a thickness of 5 μm. Specimens collected at E17.5 (data not shown), E18.5, and P0.5 were decalcified in 14% ethylenediaminetetraacetic acid (EDTA) for 7 days. Postnatal samples were decalcified for longer durations according to developmental stage. For histological analysis, sections were stained with Azan’s trichrome and imaged using an Axioscope upright light microscope (Zeiss, Germany).

Immunofluorescence staining was performed on P0.5 tissue sections. Antigen retrieval was performed in sodium citrate buffer (pH 6.0) using a pressure cooker at low pressure for 20 min. Sections were washed in PBS and blocked for 1 h at room temperature in buffer containing 2% goat serum, 5% BSA, 1% Triton X-100, and 0.1% Tween-20 in PBS. Primary antibodies diluted in blocking buffer were incubated with sections overnight at 4°C. Sections were then washed in PBS, incubated with secondary antibodies diluted in blocking buffer for 1 h, and washed again in PBS. Sections were mounted with VECTASHIELD mounting medium containing DAPI (Vector Laboratories, USA; H-1200) or counterstained with TO-PRO-3 (Invitrogen, USA; T3605) and mounted with ProLong Diamond Antifade Mountant (Invitrogen, USA; P36970). Primary antibodies used were chicken polyclonal anti-GFP (1:500; Abcam, UK; ab13970), custom-made guinea pig anti-MLL4 (1:300) (Huisman et al., 2021), rabbit monoclonal anti-SOX9 (1:300; Abcam, UK; ab185230), rabbit polyclonal anti-Ki67 (1:800; Abcam, UK; ab15580), and rabbit polyclonal anti-Cleaved Caspase-3 (1:300; Cell Signaling Technology, USA; #9661). Immunofluorescence images were acquired using an Andor Dragonfly spinning-disk confocal microscope or a THUNDER Imager fluorescence microscope (Leica Microsystems, Germany). Quantitative image analysis was performed using Fiji/ImageJ v1.54 (Schindelin et al., 2012). Individual palatal gland buds were manually delineated as regions of interest (ROIs). Within each ROI, fluorescence intensity was measured as the mean pixel intensity (arbitrary fluorescence units, AFU) in signal-positive regions identified using Analyze Particles with a minimum particle size of 10 μm^2^. Immunopositive epithelial cells within each bud were counted manually using DAPI or TO-PRO-3 counterstaining to identify individual nuclei.

### *In situ* hybridization

Multiplex fluorescent RNAscope *in situ* hybridization was performed on 5-μm-thick paraffin-embedded sections of P0.5 mouse heads using the RNAscope Multiplex Fluorescent Reagent Kit v2 (Advanced Cell Diagnostics, USA; 323100). Following deparaffinization, sections were treated with H_2_O_2_ for 10 min, subjected to target retrieval at 99°C for 13 min, and incubated with RNAscope Protease Plus at 40°C for 30 min. Mouse *Krt5* (415041-C1) and *Fgf10* (446371-C2) probes (Advanced Cell Diagnostics, USA) were hybridized at 40°C for 2 h, followed by sequential signal amplification and fluorescence detection using 520 and 570 fluorophores, respectively. Nuclei were counterstained with DAPI, and sections were mounted with ProLong Gold Antifade Mountant (Thermo Fisher Scientific, USA). *Fgf10* fluorescence intensity and the number of *Fgf10*-positive cells were quantified separately in epithelial and mesenchymal regions using Fiji/ImageJ.

Chromogenic RNAscope was performed using the RNAscope 2.5 HD Detection Reagent–BROWN kit (Advanced Cell Diagnostics, USA) according to the manufacturer’s instructions. Sections were incubated at 60°C for 1 h, deparaffinized in xylene, and treated with H_2_O_2_ for 10 min at room temperature. Target retrieval was performed at 98°C for 13 min, followed by treatment with RNAscope Protease Plus at 40°C for 30 min. Sections were hybridized with a mouse *Edar* probe (Advanced Cell Diagnostics, USA; 423011) at 40°C for 2 h, followed by signal amplification and chromogenic detection using 3,3’-diaminobenzidine (DAB). Sections were counterstained with hematoxylin, dehydrated, and mounted with VectaMount PT Permanent Mounting Medium (Vector Laboratories, USA; H-5600-60). Images were acquired using a Nikon Eclipse Ni-E microscope (Nikon Instruments, Japan). RNAscope signals were separated from hematoxylin counterstaining by color deconvolution in Fiji/ImageJ, and a uniform threshold was applied across samples to quantify the *Edar*-positive signal area.

### RNA sequencing and bioinformatics Analysis

Total RNA was extracted from microdissected soft palatal tissues using the RNeasy Micro Plus Kit (Qiagen, Germany; 74034). Four biological replicates from control mice and three from *Mll4*-cKO mice were used for sequencing, with each replicate consisting of pooled tissue from three mice. Library preparation and high-throughput sequencing were performed by Novogene (China) using the Illumina NovaSeq 6000 platform. Transcript-level expression was quantified using Salmon (Patro et al., 2017), and gene-level differential expression analysis between *Mll4*-cKO and littermate control samples was performed using DESeq2 (Love et al., 2014). Differentially expressed genes (DEGs) were defined by an absolute log2 fold change (|log2FC|) ≥ 0.4 and an adjusted *P* value ≤ 0.05.

Volcano plots were generated using EnhancedVolcano (Blighe et al., 2025). Gene Ontology Biological Process over-representation analysis (GO BP-ORA) was performed separately for downregulated and upregulated DEGs using clusterProfiler (Wu et al., 2021), with the enriched GO terms and associated genes provided in Supplementary Tables 1 and 2, respectively. A curated list of 139 salivary gland development-related genes was compiled from 15 published studies (Supplementary Table 3) and compared with the identified DEGs. Processed gene expression data are provided in the Supplementary Material.

### Statistics

Statistical analyses were performed using GraphPad Prism 10 (GraphPad Software, USA). Comparisons between control and *Mll4*-cKO groups were performed using Student’s *t*-test. Statistical significance was defined as *P* ≤ 0.05 (*), *P* ≤ 0.01 (**), and *P* ≤ 0.001 (***).

## Results

### *Mll4* deletion in the palatal mesenchyme results in palatal gland hypoplasia and ectopic cartilage formation

To investigate the role of MLL4 in palatal gland development, we generated *Mll4* conditional knockout (*Mll4*-cKO) mice by deleting *Mll4* in the palatal mesenchyme using *Osr2-Cre* (Lee et al., 2025). Around birth, when palatal gland morphogenesis begins (Hakami & Hand, 2018; Shinzato et al., 2004), *Osr2-Cre* was robustly active in the soft palate mesenchyme surrounding the developing glands, as demonstrated by the *mTmG* reporter (Figure 1A, B). Immunofluorescence confirmed efficient loss of MLL4 signal in the *Mll4*-cKO palatal mesenchyme, while MLL4 remained detectable in the adjacent epithelium (Figure 1C, D).

**Figure 1.**
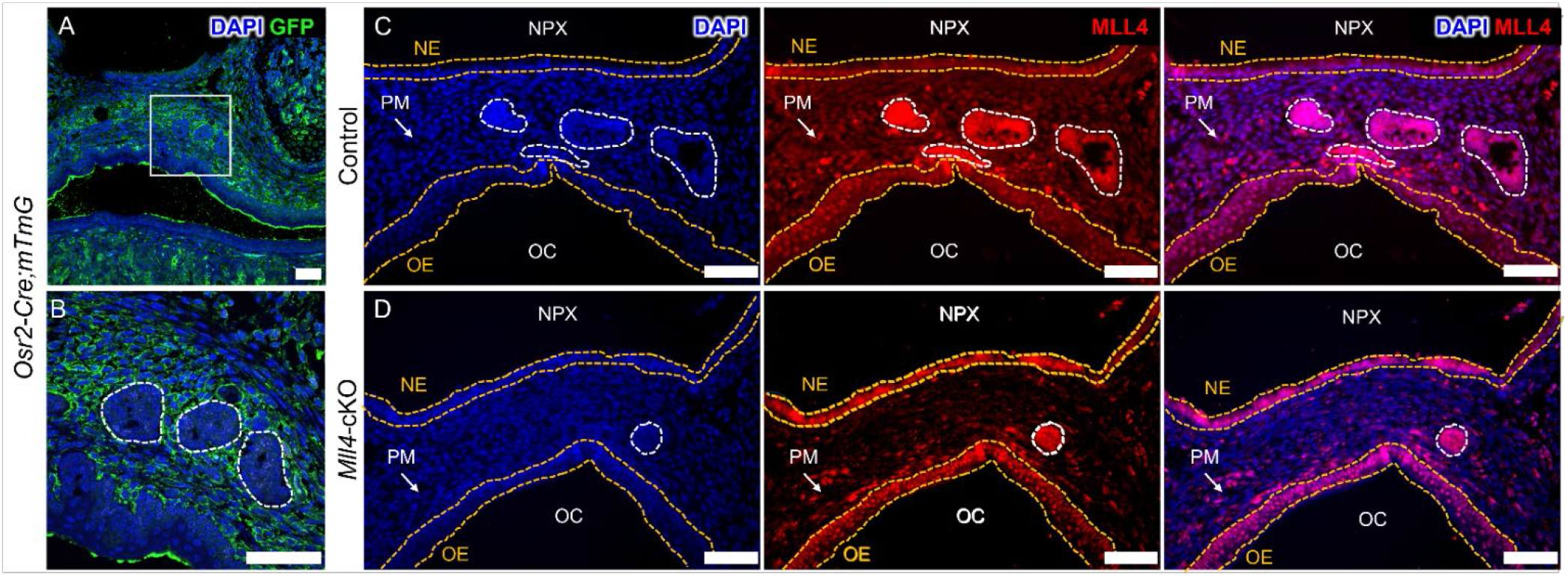
Confirmation of *Osr2-Cre* recombination and MLL4 deletion in the palatal mesenchyme at birth. (A, B) *Osr2-Cre* recombination in the soft palate at postnatal day 0.5 (P0.5), visualized by GFP (green) with DAPI counterstain (blue) in *Osr2-Cre;mT/mG* mice. (B) Higher-magnification view of the boxed region in (A). White dashed lines outline the epithelial palatal gland buds. (C, D) Immunofluorescence for MLL4 (red) with DAPI (blue) in the soft palate of control (C) and *Mll4*-cKO (D) mice at P0.5. MLL4 signal was markedly reduced in the palatal mesenchyme (PM, arrows) of *Mll4*-cKO mice, whereas MLL4 signal remained preserved in the palatal gland buds, as well as in the nasal epithelium (NE) and oral epithelium (OE). White dashed lines outline the palatal gland buds. Yellow dashed lines delineate the nasal and oral epithelial boundaries. NPX, nasopharynx; OC, oral cavity. Scale bars: 50 µm (A–D).

We next examined palatal gland morphogenesis throughout embryonic and postnatal development (Figure 2). *Mll4*-cKO mice did not develop cleft palate, allowing palatal gland development to be evaluated independently of gross alterations in the overall palatal architecture. Two prominent phenotypes were observed in *Mll4*-cKO mice: palatal gland hypoplasia and ectopic cartilage formation in the soft palate (Figure 2A–F). Palatal gland cross-sectional area was not significantly reduced at E18.5 (Figure 2A), but became significantly reduced by birth and remained reduced throughout postnatal development (Figure 2B–F). In parallel, mesenchymal condensation first became evident at E18.5 (Figure 2A), became more distinct around the developing palatal glands by P0.5 (Figure 2B), and developed into prominent ectopic cartilage from one week of age onward, exhibiting progressive tissue expansion and enlarged chondrocytes (Figure 2C–F). Collectively, these findings demonstrate that mesenchymal deletion of *Mll4* results in persistent palatal gland hypoplasia accompanied by ectopic cartilage formation within the developing soft palate.

**Figure 2.**
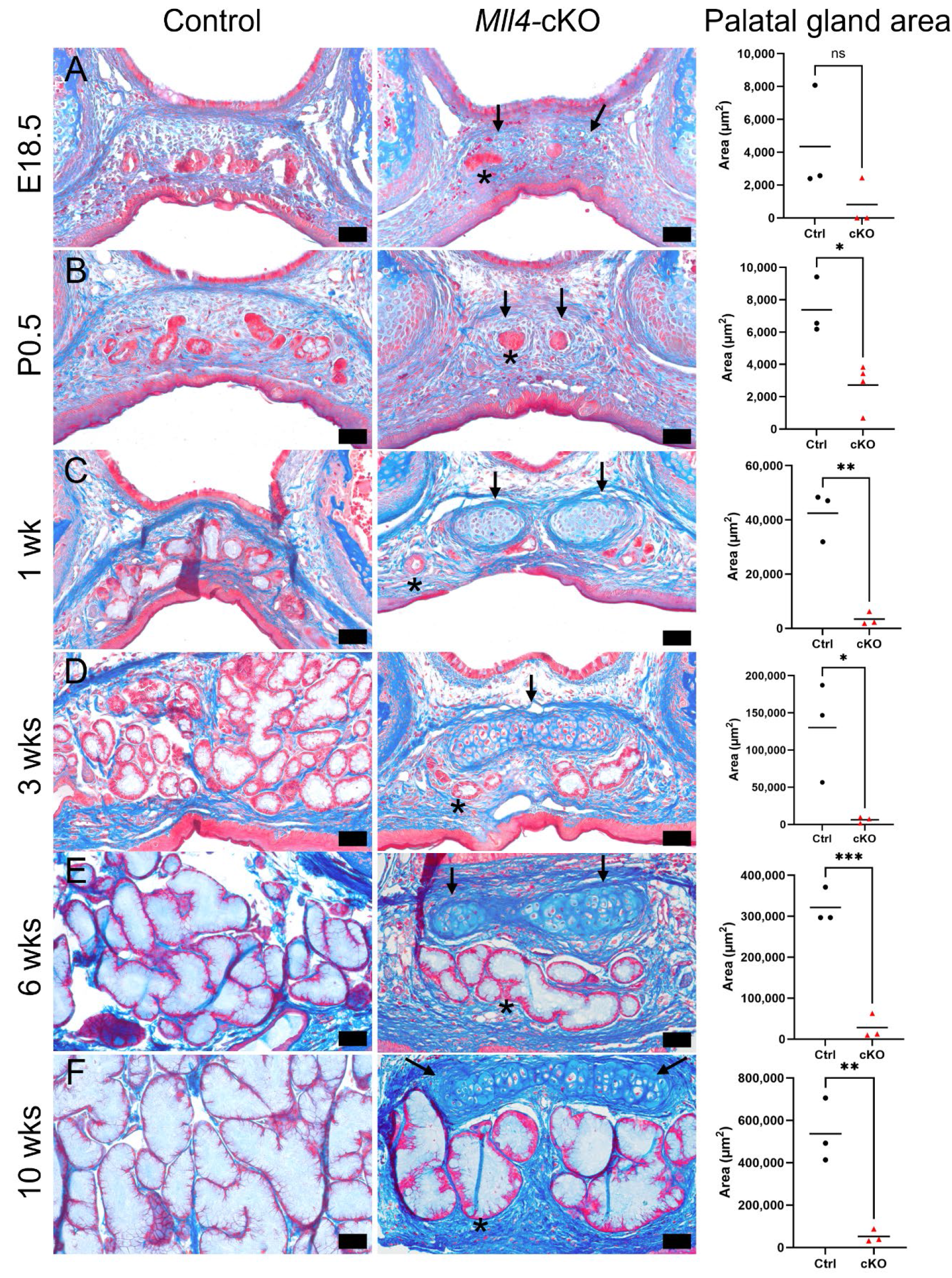
*Mll4* deficiency induces ectopic cartilage formation and palatal gland hypoplasia in the soft palate. (A–F) Azan’s trichrome staining of the middle region of the soft palate at different developmental stages in control and *Mll4*-cKO mice: E18.5 (A), P0.5 (B), 1 week (C), 3 weeks (D), 6 weeks (E), and 10 weeks (F). Arrows indicate ectopic cartilage, and asterisks indicate hypoplastic palatal glands. Right panels show quantification of total palatal gland cross-sectional area, measured by manually delineating individual palatal gland buds using FIJI software and summing the cross-sectional areas of all palatal gland buds within each section. *n* = 3 mice per genotype at each stage. Statistical analysis was performed using Student’s *t*-test (ns, not significant; \**P* < 0.05; \*\**P* < 0.01; \*\*\**P* < 0.001). Scale bars: 50 µm (A–F).

### *Mll4* deletion alters transcriptional programs associated with palatal gland development and chondrogenesis

To investigate the molecular basis of the observed phenotypes in *Mll4*-cKO mice, we performed RNA-seq on microdissected soft palate tissues from control and *Mll4*-cKO mice at P0.5, when the phenotype first became evident. We identified 789 differentially expressed genes (DEGs), including 268 downregulated and 521 upregulated genes, using an adjusted *P* ≤ 0.05 and |log2 fold change| ≥ 0.4 (Figure 3A). Gene Ontology Biological Process over-representation analysis (GO BP-ORA) identified enriched biological processes related to salivary gland development, branching morphogenesis, FGF/WNT signaling, cartilage development, and chondrocyte differentiation among downregulated genes (Figure 3B; Supplementary Table 1). In contrast, upregulated genes were enriched for biological processes related to muscle differentiation, motile cilium assembly, chondrocyte proliferation, and positive regulation of chondrocyte proliferation (Figure 3B′; Supplementary Table 2). Notably, GO terms associated with salivary gland morphogenesis and cartilage differentiation were enriched among downregulated genes, whereas those associated with chondrocyte proliferation were enriched among upregulated genes in *Mll4*-cKO mice.

**Figure 3.**
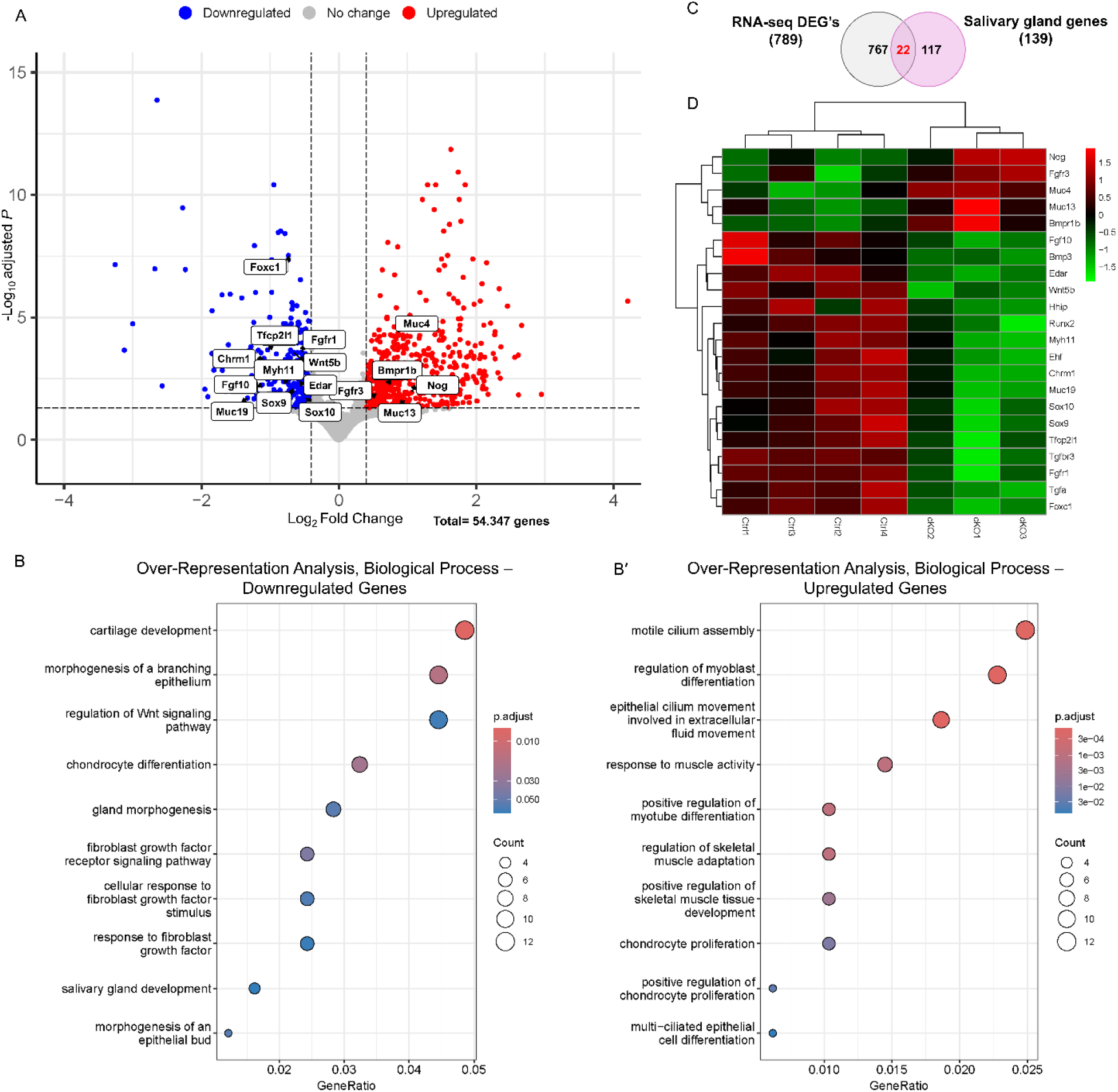
*Mll4* loss disrupts key transcriptional networks governing salivary gland development and chondrocyte proliferation. (A–D) Genome-wide mRNA expression analysis was performed using microdissected soft palate tissues from control and *Mll4*-cKO mice at birth (P0.5). (A) Volcano plot showing that 789 genes were differentially expressed (adjusted *P* ≤ 0.05, |log2 fold change| ≥ 0.4): 268 downregulated and 521 upregulated. (B, B′) Gene Ontology Biological Process over-representation analysis (GO BP-ORA) of the differentially expressed genes (DEGs). GO terms enriched among downregulated genes (B) were associated with salivary gland development, branching morphogenesis, cartilage development, and FGF/WNT signaling (Supplementary Table 1). GO terms enriched among upregulated genes (B′) were associated with muscle differentiation, motile cilium assembly, and chondrocyte proliferation (Supplementary Table 2). (C) Venn diagram showing the overlap between the 789 DEGs and 139 genes previously implicated in salivary gland development, identifying 22 overlapping genes. (D) Heatmap showing the expression patterns of the 22 overlapping genes identified in (C): 17 downregulated and 5 upregulated in *Mll4*-cKO mice.

To further prioritize candidate regulators of palatal gland development, we compared the DEGs with 139 genes previously implicated in salivary gland development (Chibly et al., 2022; Das et al., 2009; Emmerson & Knox, 2018; Gluck et al., 2021; Hand et al., 2024; Hauser et al., 2020; Langthasa et al., 2025; Mattingly et al., 2015; Rose et al., 2023; Sisto et al., 2022; Song et al., 2018; Suzuki et al., 2021; Teshima et al., 2011; Yamaguchi et al., 2006; Yamamoto et al., 2008) (Supplementary Table 3). This comparison identified 22 overlapping genes: 17 downregulated and 5 upregulated in *Mll4*-cKO mice (Figure 3C, D). The downregulated genes included *Fgf10*, *Sox9*, *Sox10*, *Edar*, *Wnt5b*, *Foxc1*, *Tfcp2l1*, *Fgfr1*, *Muc19,* and *Runx2*, whereas the upregulated genes included *Nog*, *Bmpr1b*, and *Fgfr3*. Based on their established roles in epithelial–mesenchymal signaling during salivary gland development, *Fgf10*, *Sox9*, and *Edar* were selected for subsequent validation.

### *Mll4* deletion reduces mesenchymal *Fgf10* and epithelial SOX9 and *Edar* expression

To validate the RNA-seq findings, we next examined the expression of *Fgf10*, SOX9, and *Edar* in the developing soft palate at P0.5. Multiplex RNAscope *in situ* hybridization revealed a significant reduction in *Fgf10* expression in the palatal mesenchyme surrounding the developing palatal glands of *Mll4*-cKO mice (Figure 4A, B). In contrast, *Fgf10* expression in the palatal gland epithelium was not significantly altered (Figure 4C). Quantitative analysis further demonstrated significant reductions in both *Fgf10* expression and the number of *Fgf10*-positive mesenchymal cells (Figure 4D), confirming selective loss of mesenchymal *Fgf10* expression following *Mll4* deletion.

**Figure 4.**
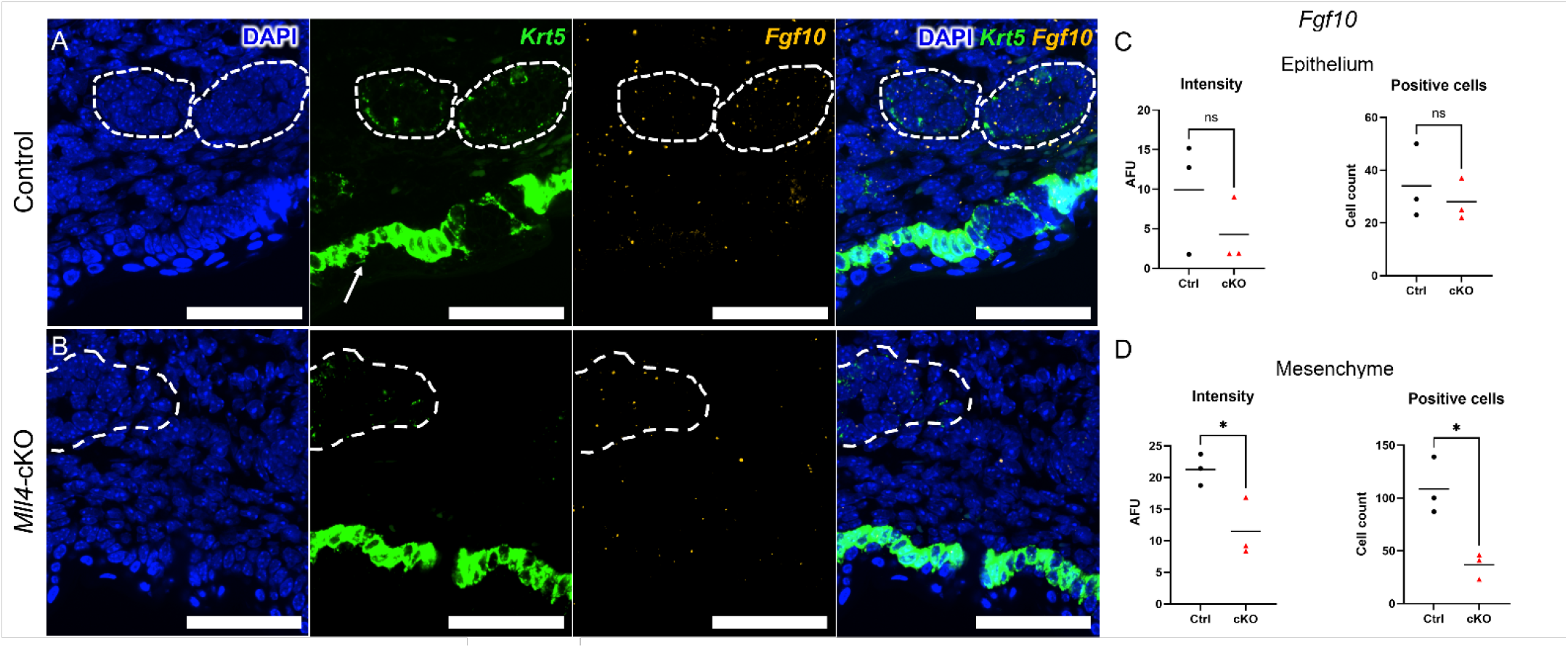
Reduced expression of mesenchymal *Fgf10* in the *Mll4*-cKO soft palate. (A, B) Multiplex RNAscope *in situ* hybridization in control and *Mll4*-cKO mice at postnatal day 0.5 (P0.5), showing *Fgf10* expression (yellow) within the soft palate region. The epithelial marker *Krt5* (green) delineates the palatal gland buds (dashed outlines); pronounced *Krt5* expression is also observed in the oral epithelium (arrows). Nuclei are counterstained with DAPI (blue). (C) Quantification of *Fgf10* signal intensity (arbitrary fluorescence units, AFU) and the number of *Fgf10*-positive cells within the epithelial compartment, showing no significant differences between genotypes. (D) Quantification of *Fgf10* signal intensity and the number of *Fgf10*-positive cells within the mesenchymal compartment, showing significant reductions in both parameters in *Mll4*-cKO mice compared to controls. *n* = 3 mice per genotype. Statistical analysis was performed using Student’s *t*-test (ns, not significant; \**P* < 0.05). Scale bars: 50 µm (A, B).

We next examined SOX9 protein expression by immunofluorescence. Consistent with the RNA-seq results, SOX9 expression was markedly reduced in the palatal gland epithelium of *Mll4*-cKO mice (Figure 5A, B). In contrast, strong SOX9 immunoreactivity was detected within the ectopic cartilage that developed adjacent to the palatal glands in *Mll4*-cKO mice (Figure 5A, B). Quantitative analysis confirmed significant reductions in both SOX9 fluorescence intensity and the number of SOX9-positive epithelial cells (Figure 5C, D).

**Figure 5.**
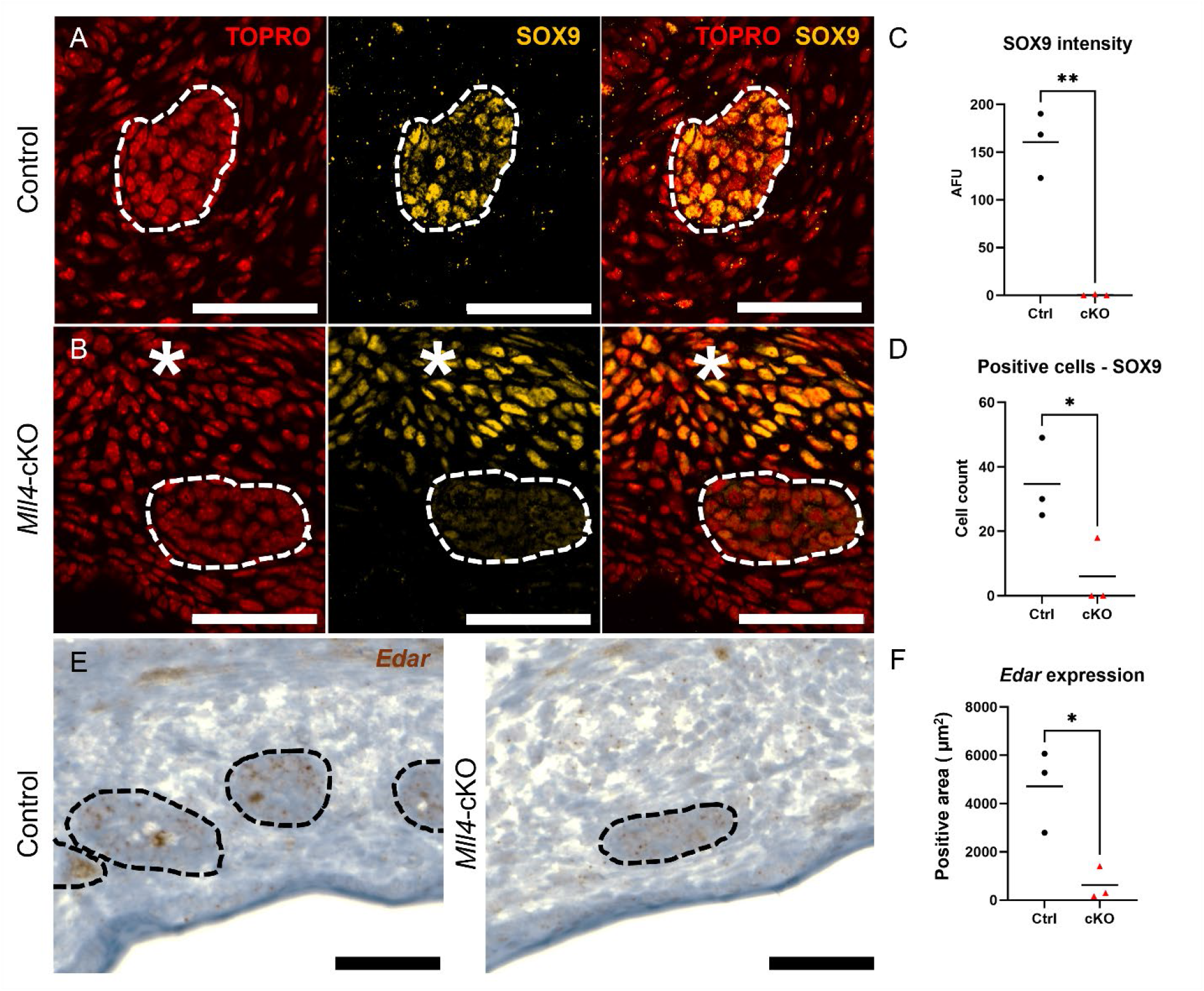
Reduced expression of epithelial SOX9 and *Edar* in the *Mll4-*cKO soft palate. (A, B) Immunofluorescence for SOX9 (yellow) with TO-PRO-3 nuclear counterstain (red) in the soft palate of control (A) and *Mll4*-cKO (B) mice at postnatal day 0.5 (P0.5). Dashed lines delineate the epithelial palatal gland buds. The asterisk indicates adjacent ectopic cartilage in *Mll4*-cKO mice. SOX9 expression was markedly reduced in the palatal gland buds and increased within the adjacent ectopic cartilage of *Mll4*-cKO mice. (C) Quantification of SOX9 signal intensity (AFU) within the palatal gland buds. (D) Quantification of the number of SOX9-positive epithelial cells within the palatal gland buds. (E) RNAscope *in situ* hybridization for *Edar* expression (brown) in the palatal gland buds (dashed outlines) of control and *Mll4*-cKO mice at P0.5. (F) Quantification of the total *Edar*-positive hybridization area per field. *n* = 3 mice per genotype. Statistical analysis was performed using Student’s *t*-test (\**P* < 0.05; \*\**P* < 0.01). Scale bars: 50 µm (A, B, E).

Finally, RNAscope *in situ* hybridization demonstrated a significant reduction in *Edar* expression in the palatal gland epithelium of *Mll4*-cKO mice (Figure 5E, F). Together, these findings validate the transcriptomic results and demonstrate that mesenchymal *Mll4* deletion is associated with reduced mesenchymal *Fgf10* expression accompanied by reduced epithelial SOX9 and *Edar* expression.

### *Mll4* deletion reduces epithelial cell proliferation in the developing palatal gland

Because *Fgf10* and SOX9 are key regulators of epithelial progenitor expansion during salivary gland development, we next examined cell proliferation and apoptosis in the developing palatal gland at P0.5. Immunofluorescence for Ki67 revealed a significant reduction in the number of proliferating epithelial cells within the palatal gland buds of *Mll4*-cKO mice compared to controls (Figure 6A–C). In contrast, apoptosis, assessed by cleaved caspase-3 immunostaining, was not significantly different between genotypes (data not shown). These findings indicate that palatal gland hypoplasia in *Mll4*-cKO mice is associated with reduced epithelial proliferation, without a detectable increase in apoptosis at P0.5.

**Figure 6.**
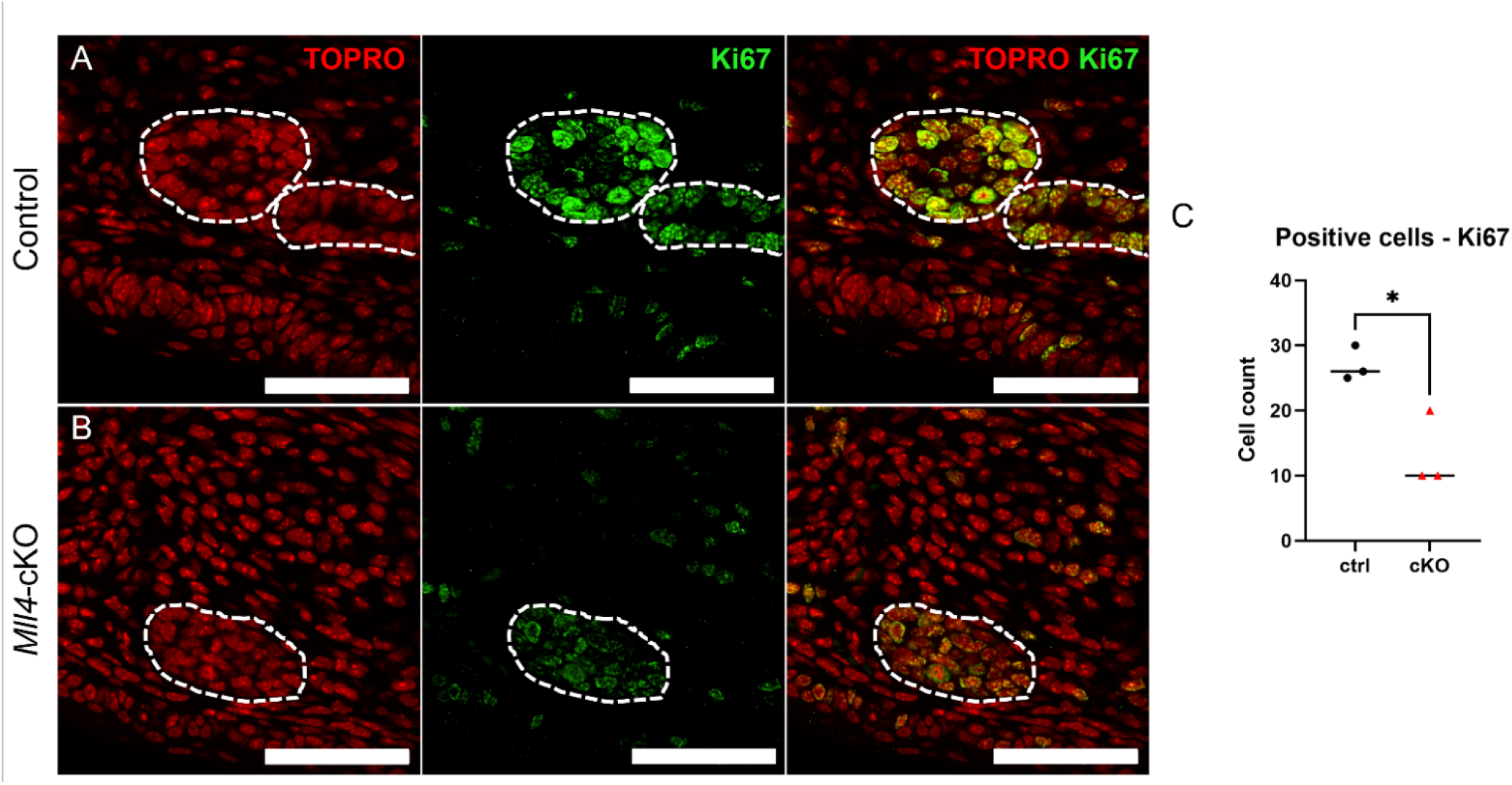
**Reduced epithelial cell proliferation in the *Mll4*-cKO palatal gland**. (A, B) Immunofluorescence for the proliferation marker Ki67 (green) with TO-PRO-3 nuclear counterstain (red) in the palatal gland of control (A) and *Mll4*-cKO (B) mice at postnatal day 0.5 (P0.5). Dashed lines delineate the epithelial palatal gland buds. Ki67-positive epithelial cells were markedly reduced in *Mll4*-cKO mice. (C) Quantification of the number of Ki67-positive epithelial cells within the palatal gland buds. *n* = 3 mice per genotype. Statistical analysis was performed using Student’s *t*-test (\**P* < 0.05). Scale bars: 50 µm (A, B).

## Discussion

In this study, we identified MLL4 as an essential epigenetic regulator of palatal gland development. Palatal mesenchyme-specific deletion of *Mll4* resulted in palatal gland hypoplasia accompanied by ectopic cartilage formation in the soft palate, revealing an unexpected requirement for mesenchymal MLL4 in coordinating normal gland morphogenesis. Transcriptomic profiling, together with molecular and cellular validation, demonstrated that loss of mesenchymal MLL4 was associated with reduced mesenchymal *Fgf10* expression, decreased epithelial SOX9 and *Edar* expression, and impaired epithelial proliferation within the developing palatal gland. Because *Mll4* deletion was restricted to the palatal mesenchyme, whereas the principal molecular and cellular defects occurred within the epithelial palatal gland buds, our findings support a non-cell-autonomous mechanism in which mesenchymal MLL4 may maintain *Fgf10* expression and thereby contribute to epithelial proliferation and palatal gland morphogenesis.

### MLL4 regulates epithelial–mesenchymal signaling during palatal gland development

Our findings suggest that mesenchymal MLL4 regulates palatal gland development through epithelial– mesenchymal signaling. Loss of mesenchymal *Mll4* reduced *Fgf10* expression in the palatal mesenchyme, accompanied by decreased epithelial SOX9 and *Edar* expression and reduced epithelial proliferation. This pattern is consistent with the well-established mesenchymal FGF10–epithelial FGFR2b signaling axis that governs major salivary gland development. Mesenchyme-derived FGF10 signals through epithelial FGFR2b to promote epithelial progenitor proliferation and branching morphogenesis, whereas disruption of this pathway results in gland hypoplasia or aplasia and can arrest salivary gland development near the initial bud stage (Jaskoll et al., 2005; Steinberg et al., 2005). Downstream of FGF10, SOX9 sustains the identity and proliferative capacity of distal epithelial progenitors, and its deletion impairs epithelial proliferation and branching morphogenesis (Chatzeli et al., 2017). Together, these findings suggest that mesenchymal MLL4 maintains palatal gland morphogenesis by preserving this core epithelial–mesenchymal signaling pathway.

Importantly, the epithelial abnormalities observed in the palatal gland buds of *Mll4*-cKO mice are unlikely to result from a cell-autonomous requirement for MLL4 within the epithelial palatal gland buds. Immunofluorescence confirmed that MLL4 remained readily detectable in the palatal gland buds of *Mll4*-cKO mice despite efficient deletion in the surrounding palatal mesenchyme (Figure 1C, D). Nevertheless, SOX9 and *Edar* expression, together with epithelial proliferation, were all significantly reduced in the palatal gland buds (Figures 5, 6). Taken together, these findings suggest that the epithelial defects may arise indirectly from altered mesenchymal FGF10 signaling rather than from loss of epithelial MLL4 itself. Among the mesenchymal regulators identified in our transcriptomic analysis, *Fgf10* emerged as the strongest candidate mediator because its expression was selectively reduced in the palatal mesenchyme while remaining unchanged within the epithelial compartment (Figure 4). Notably, *Fgfr2* expression was not significantly altered in the RNA-seq dataset, suggesting that reduced mesenchymal FGF10 availability, rather than decreased Fgfr2 expression, may contribute to the epithelial defects observed in *Mll4*-cKO mice.

In addition to reduced mesenchymal *Fgf10* expression, our transcriptomic analysis identified other differentially expressed developmental regulators that may contribute to the palatal gland defects observed in *Mll4*-cKO mice . Among these, *Sox10* was significantly downregulated. SOX10 functions downstream of SOX9 during lacrimal gland development (Chen et al., 2014) and is required for the formation and maintenance of secretory units, with epithelial deletion of *Sox10* resulting in marked loss of secretory acini while largely preserving the ductal tree (Athwal et al., 2019). Furthermore, FGF signaling has been shown to activate a SOX9–SOX10 transcriptional program during lacrimal gland branching morphogenesis (Chen et al., 2014), suggesting that reduced *Sox10* expression may reflect impaired epithelial differentiation secondary to decreased mesenchymal FGF10 signaling.

*Edar* was also significantly downregulated in *Mll4*-cKO tissue. During major salivary gland development, EDA is produced predominantly by the mesenchyme, where its expression is induced by canonical WNT/β-catenin signaling, whereas EDAR is primarily expressed in the glandular epithelium and promotes epithelial branching morphogenesis through NF-κB signaling (Häärä et al., 2011). Because *Eda* expression was not significantly altered in our RNA-seq dataset, the contribution of altered EDA ligand availability to reduced epithelial *Edar* expression remains unclear. Decreased *Edar* expression may reflect impaired epithelial progenitor maintenance or delayed epithelial morphogenesis associated with reduced mesenchymal FGF10 signaling and diminished epithelial growth. Alternatively, reduced *Edar* expression may be a consequence of the hypoplastic epithelial compartment observed in *Mll4*-cKO mice.

Additional developmental regulators identified in the RNA-seq dataset included *Wnt5b*, *Tfcp2l1*, and *Foxc1*. Previous studies have implicated WNT5B-mediated non-canonical WNT signaling in major salivary gland ductal differentiation through regulation of TFCP2L1 (Mattingly et al., 2015; Patel et al., 2011). FOXC1 has also been implicated in salivary gland organoid formation, where it cooperates with SOX9 to promote salivary gland rudiment formation (Tanaka et al., 2018). Furthermore, studies in the mouse lacrimal gland have demonstrated that FOXC1 mediates mesenchymal responses to BMP signaling during branching morphogenesis (Mattiske et al., 2006). Consistent with the involvement of BMP-associated developmental pathways, several genes, including *Bmpr1b*, *Nog*, and *Bmp3*, were differentially expressed in *Mll4*-cKO tissue. Because both positive and negative regulators of BMP signaling were altered, these findings suggest broader perturbation of BMP-associated developmental programs without establishing whether overall BMP signaling is increased or decreased.

Finally, *Muc19*, which encodes a major secreted mucin produced by mature mucous acinar cells (Das et al., 2009), was markedly downregulated. Unlike the membrane-associated mucins *Muc13* and *Muc4*, which were upregulated, reduced *Muc19* expression is consistent with impaired maturation and secretory differentiation of the developing palatal glands. Collectively, these findings suggest that MLL4-dependent regulation of palatal gland development extends beyond the FGF10 signaling axis to encompass additional pathways involved in epithelial morphogenesis and gland maturation.

### Ectopic cartilage formation in the *Mll4*-cKO soft palate

An unexpected finding of this study was the development of ectopic cartilage within the soft palate following mesenchymal deletion of *Mll4*. Consistent with this phenotype, GO analysis revealed a distinct pattern of dysregulated chondrogenic programs. Biological processes related to cartilage development and chondrocyte differentiation were significantly enriched among downregulated genes, including *Sox9*, *Wnt5b*, *Ltbp3*, *Runx2*, *Bmp3*, and *Fgfr1* (Figure 3B; Supplementary Table 1). In contrast, biological processes associated with chondrocyte proliferation and its positive regulation were enriched among upregulated genes, including *Pbxip1*, *Ccn3*, *Bmpr1b*, *Ltf*, and *Mustn1* (Figure 3B; Supplementary Table 2). At the protein level, SOX9 expression was markedly reduced in the epithelial palatal gland buds but strongly increased within the ectopic cartilage (Figure 5A–D). Together, these findings suggest that mesenchymal *Mll4* deletion does not simply enhance or suppress cartilage formation but rather disrupts the balance between chondrocyte differentiation and proliferative programs during soft palate development.

The transcriptomic changes further support this interpretation. Although *Runx2*, a key regulator of osteogenic differentiation and chondrocyte maturation, was reduced in the bulk RNA-seq dataset, *Acan* expression was increased, whereas *Col2a1* and *Col10a1* were not significantly altered. This expression pattern, together with the localized accumulation of SOX9-positive cells within the ectopic cartilage, is compatible with the emergence of a focal cartilage-producing population without clear evidence of progression toward a normal hypertrophic differentiation program. Because RNA-seq was performed using whole soft palate tissue, however, these transcriptional changes likely reflect the overall developmental state of the tissue rather than the molecular profile of the ectopic cartilage itself.

Although the mechanism underlying ectopic cartilage formation in *Mll4*-cKO mice remains unclear, the combined histological and transcriptomic findings suggest that mesenchymal MLL4 regulates developmental programs required for proper craniofacial mesenchymal patterning. Because the ectopic cartilage occupies only a limited region of the soft palate, whereas the RNA-seq dataset represents the entire dissected tissue, the apparent discrepancy between localized ectopic cartilage formation and reduced expression of several cartilage developmental regulators is not unexpected. Rather, these data suggest that loss of mesenchymal MLL4 results in abnormal spatial regulation of chondrogenic differentiation rather than a uniform increase in cartilage formation throughout the soft palate. Future lineage-tracing and spatial transcriptomic studies will therefore be required to determine the cellular origin of the ectopic cartilage and to distinguish direct effects of MLL4 loss from secondary changes in the surrounding tissue.

Interestingly, the ectopic cartilage phenotype observed in the present study contrasts with our previous findings in the midpalatal suture using the same *Mll4^fl/fl^;Osr2-Cre* mouse model, in which mesenchymal *Mll4* deletion impaired cartilage formation and reduced the expression of multiple chondrogenic markers, including SOX9, *Acan*, and *Col2a1* (Lee et al., 2025). By contrast, *Mll4*-cKO mice in the present study developed ectopic cartilage with increased local SOX9 expression, accompanied by upregulation of chondrocyte proliferation-related programs in the transcriptomic analysis. These contrasting phenotypes suggest that the function of MLL4 in craniofacial mesenchyme is highly dependent on developmental context. Accordingly, MLL4 may differentially regulate skeletal lineage programs according to the local signaling environment and tissue-specific developmental cues.

### Clinical implications of MLL4-dependent palatal gland development

Pathogenic variants in KMT2D, the gene encoding MLL4, cause Kabuki syndrome, a congenital disorder characterized by distinctive craniofacial abnormalities and developmental defects (Boniel et al., 2021). Although craniofacial and oral manifestations of Kabuki syndrome are well recognized, salivary gland abnormalities have not been reported as a characteristic feature of the disorder. Our findings therefore identify a previously unrecognized developmental function of MLL4 in palatal gland morphogenesis and provide a framework for investigating the role of KMT2D in salivary gland development and its potential contribution to the craniofacial manifestations of KMT2D deficiency.

More broadly, our findings implicate mesenchymal MLL4–FGF10 signaling in the epithelial– mesenchymal interactions required for palatal gland development. These findings may also inform future studies of congenital salivary gland disorders and regenerative approaches aimed at restoring gland morphogenesis by re-establishing appropriate epithelial–mesenchymal signaling.

### Limitations and future directions

Several limitations of this study should be acknowledged. First, our transcriptomic analysis was based on bulk RNA-seq of microdissected soft palate tissue, limiting resolution of cell type-specific transcriptional changes. Future single-cell transcriptomic approaches will be required to define the epithelial and mesenchymal contributions of the candidate genes identified in this study. Second, although our findings support an important regulatory role for MLL4, we did not determine its direct genomic targets. Future chromatin profiling studies, such as CUT&RUN or ChIP-seq, will therefore be important for defining the transcriptional network controlled by MLL4 in the palatal mesenchyme. Finally, the cellular origin of the ectopic cartilage remains unresolved. Lineage-tracing studies will be required to determine whether ectopic chondrogenesis results from altered mesenchymal cell fate or other developmental mechanisms. Addressing these questions will further advance our understanding of how MLL4 coordinates epithelial–mesenchymal signaling during palatal gland development.

## Supporting information

Supplementary RNA-seq Data (All Genes)

## Statements

### Data Availability Statement

Processed gene expression data and differential expression analysis results generated in this study are provided in the Supplementary Material. The original RNA-sequencing read files are no longer available due to data loss and therefore cannot be deposited in a public sequencing repository. Further inquiries regarding the available processed data may be directed to the corresponding authors.

### Ethics Statement

The animal study was reviewed and approved by Institutional Animal Care and Use Committee (IACUC) at the University at Buffalo.

### Author contributions

BP: Data curation, Formal analysis, Investigation, Methodology, Visualization, Writing – original draft, Writing – review and editing. HJ: Investigation, Methodology, Data curation, Visualization, Writing – review and editing. YP: Data curation, Formal analysis, Writing – review and editing. R-AR: Resources, Supervision, Writing – review and editing. S-KL: Resources, Writing – review and editing. JWL: Resources, Writing – review and editing. J-ML: Conceptualization, Formal analysis, Investigation, Methodology, Funding acquisition, Supervision, Validation, Writing – review and editing. H-JEK: Conceptualization, Formal analysis, Funding acquisition, Project administration, Resources, Supervision, Validation, Writing – original draft, Writing – review and editing.

## Acknowledgments

All fluorescence imaging was performed at the Optical Imaging and Analysis Facility (OIAF), School of Dental Medicine, University at Buffalo, The State University of New York. We thank Andrew D. McCall at the OIAF for his assistance with fluorescence microscopy and image analysis. We also thank Jason Osinski in the Romano laboratory for helpful guidance on immunofluorescence protocols. Finally, we thank the members of the Kwon laboratory and our colleagues in the Department of Oral Biology for their helpful comments and feedback.

## Generative AI statement

The authors declare that generative AI tools, including Google Gemini, OpenAI ChatGPT, and Anthropic Claude, were used to assist with language editing, clarity, and manuscript refinement. All AI-assisted content was critically reviewed and verified by the authors, who take full responsibility for the accuracy and integrity of the manuscript.

