## Supplemental Data 1 for "MLL4 regulates palatal salivary gland development"

### SUPPLEMENTAL APPENDIX

| Over Representation Analysis (ORA) Biological Process – Downregulated Genes | | |
|---|---|---|
| **GO ID** | **Description** | **Gene ID** |
| GO:0051216 | cartilage development | *Sox9/Wnt5b/Ltbp3/Runx2/Rspo2/Creb3l2/Bmp3/Thra/Barx2/Osr2/Chadl/Fgfr1* |
| GO:0061138 | morphogenesis of a branching epithelium | *Sox9/Hhip/Rspo2/Rdh10/Sema5a/Agt/Fgf10/Slit2/Ar/Sox10/Fgfr1* |
| GO:0030111 | regulation of Wnt signaling pathway | *Sox9/Wnt5b/Sox13/Rspo2/Rspo1/Nkd2/Sema5a/Thra/Fgf10/Klf15/Sox10* |
| GO:0002062 | chondrocyte differentiation | *Sox9/Wnt5b/Ltbp3/Runx2/Creb3l2/Osr2/Chadl/Fgfr1* |
| GO:0022612 | gland morphogenesis | *Id4/Sox9/Edar/Fgf10/Slit2/Ar/Fgfr1* |
| GO:0008543 | fibroblast growth factor receptor signaling pathway | *Hhip/Creb3l1/Runx2/Fgf10/Flrt2/Fgfr1* |
| GO:0044344 | cellular response to fibroblast growth factor stimulus | *Hhip/Creb3l1/Runx2/Fgf10/Flrt2/Fgfr1* |
| GO:0071774 | response to fibroblast growth factor | *Hhip/Creb3l1/Runx2/Fgf10/Flrt2/Fgfr1* |
| GO:0007431 | salivary gland development | *Tfcp2l1/Edar/Fgf10/Fgfr1* |
| GO:0060572 | morphogenesis of an epithelial bud | *Rdh10/Fgf10/Ar* |

**Supplementary Table 1. Enriched GO biological process terms for downregulated genes.** Over-representation analysis (ORA) showing significantly enriched Gene Ontology (GO) biological process terms among genes downregulated in *Mll4*-cKO mice compared with controls.

| Over Representation Analysis (ORA) Biological Process – Upregulated Genes | | |
|---|---|---|
| **GO ID** | **Description** | **Gene ID** |
| GO:0044458 | motile cilium assembly | *Ccdc181/Foxj1/Spag17/Cep128/Cfap58/Cfap119/Ttc12/Cfap69/Dnali1/Lrrc46/Ccdc40/Iqcg* |
| GO:0045661 | regulation of myoblast differentiation | *Mef2c/Capn3/Csrp3/Myf6/Mstn/Myog/Klhl41/Rbm24/Ankrd2/Mustn1/Smyd1* |
| GO:0003351 | epithelial cilium movement involved in extracellular fluid movement | *Rsph14/Ak7/Kif27/Spag17/Nme5/Cfap45/Aqp4/Rsph1/Ccdc40* |
| GO:0014850 | response to muscle activity | *Prkag3/Srl/Capn3/Ryr2/Fndc5/Mstn/Myog* |
| GO:0010831 | positive regulation of myotube differentiation | *Tbx1/Csrp3/Myog/Rbm24/Smyd1* |
| GO:0014733 | regulation of skeletal muscle adaptation | *Tnnc1/Tnni1/Cmya5/Tnnt1/Mymk* |
| GO:0048643 | positive regulation of skeletal muscle tissue development | *Usp2/Mef2c/Lmod3/Myf6/Myog* |
| GO:0035988 | chondrocyte proliferation | *Pbxip1/Ccn3/Bmpr1b/Ltf/Mustn1* |
| GO:1902732 | positive regulation of chondrocyte proliferation | *Pbxip1/Ltf/Mustn1* |
| GO:1903251 | multi-ciliated epithelial cell differentiation | *Foxj1/Wfdc2/Deup1* |

**Supplementary Table 2. Enriched GO biological process terms for upregulated genes.** Over-representation analysis (ORA) showing significantly enriched Gene Ontology (GO) biological process terms among genes upregulated in *Mll4*-cKO mice compared with controls.

| | | | | |
|---|---|---|---|---|
| *Acta2* | *Edaradd* | *Grem1* | *Myh11* | *Shh* |
| *Acvr2b* | *Egf* | *Grhl2* | *Myl9* | *Six1* |
| *Aldh1a2* | *Egfr* | *Gsk3b* | *Nfib* | *Slc12a2* |
| *Amy1* | *Ehf* | *Gstt1* | *Nfkb1* | *Smgc* |
| *Aqp5* | *Elf5* | *Hbegf* | *Nkx2-3* | *Smo* |
| *Artn* | *Erbb2* | *Hhip* | *Nkx3-1* | *Sox10* |
| *Ascl3* | *Erbb3* | *Hif1a* | *Nog* | *Sox2* |
| *Bhlha15* | *Erbb4* | *Ihh* | *Notch1* | *Sox9* |
| *Bmp1* | *Fgf1* | *Irf6* | *Notch2* | *Spry1* |
| *Bmp2* | *Fgf10* | *Jag1* | *Notch3* | *Spry2* |
| *Bmp3* | *Fgf13* | *Jag2* | *Notch4* | *T* |
| *Bmp4* | *Fgf2* | *Kif3a* | *Nrg1* | *Tfcp2l1* |
| *Bmp6* | *Fgf3* | *Kit* | *Nrtn* | *Tgfa* |
| *Bmp7* | *Fgf7* | *Krt14* | *Pax6* | *Tgfb1* |
| *Bmpr1a* | *Fgf8* | *Krt18* | *Pax9* | *Tgfb2* |
| *Bmpr1b* | *Fgfr1* | *Krt19* | *Pdgfa* | *Tgfb3* |
| *Bmpr2* | *Fgfr2* | *Krt5* | *Pdgfra* | *Tgfbr1* |
| *Cdh1* | *Fgfr3* | *Krt7* | *Pdgfrb* | *Tgfbr2* |
| *Cftr* | *Fgfr4* | *Krt8* | *Pik3ca* | *Tgfbr3* |
| *Chrm1* | *Foxa1* | *Lgr4* | *Pitx1* | *Tnf* |
| *Chrm3* | *Foxc1* | *Lgr5* | *Prol1* | *Trp63* |
| *Creb1* | *Gdnf* | *Lgr6* | *Prom1* | *Vip* |
| *Dhh* | *Gfra1* | *Mapk1* | *Pspn* | *Wnt1* |
| *Dlk1* | *Gfra2* | *Mapk3* | *Ptch1* | *Wnt3a* |
| *Dlk2* | *Gfra3* | *Muc13* | *Ptch2* | *Wnt5a* |
| *Dll1* | *Gli1* | *Muc19* | *Ret* | *Wnt5b* |
| *Eda* | *Gli2* | *Muc4* | *Runx1* | *Yap1* |
| *Edar* | *Gli3* | *Myc* | *Runx2* | |

**Supplementary Table 3. List of the 139 salivary gland development-related genes.** A total of 139 genes were compiled from previous studies (Chibly et al., 2022; Das et al., 2009; Emmerson & Knox, 2018; Gluck et al., 2021; Hand et al., 2024; Hauser et al., 2020; Langthasa et al., 2025; Mattingly et al., 2015; Rose et al., 2023; Sisto et al., 2022; Song et al., 2018; Suzuki et al., 2021; Teshima et al., 2011; Yamaguchi et al., 2006; Yamamoto et al., 2008).
